# Targeting the CALCB/RAMP1-axis inhibits growth of Ewing sarcoma

**DOI:** 10.1101/491100

**Authors:** Marlene Dallmayer, Jing Li, Shunya Ohmura, Rebeca Alba-Rubio, Michaela C. Baldauf, Tilman L. B. Hölting, Julian Musa, Max M. L. Knott, Stefanie Stein, Florencia Cidre-Aranaz, Fabienne S. Wehweck, Laura Romero-Pérez, Julia S. Gerke, Martin F. Orth, Aruna Marchetto, Thomas Kirchner, Horacio Bach, Giuseppina Sannino, Thomas G. P. Grünewald

**Affiliations:** Max-Eder Research Group for Pediatric Sarcoma Biology, Institute of Pathology of the LMU Munich, Munich, Germany; Institute of Pathology of the LMU Munich, Munich, Germany; German Cancer Consortium (DKTK), Heidelberg, Germany; German Cancer Research Center (DKFZ), Heidelberg, Germany; Department of Medicine, Division of Infectious Diseases, and IIRC Antibody Engineering and Proteomics facility, University of British Columbia, Vancouver, BC, Canada

**Author notes:** **Corresponding author**: Thomas G. P. Grünewald, MD, PhD, Max-Eder Research Group for Pediatric Sarcoma Biology, Institute of Pathology of the LMU Munich, Thalkirchner Str. 36, 80337 Munich, Germany, Web www.lmu.de/sarkombiologie.

**Keywords:** Ewing sarcoma, CALCB, CGRP2, RAMP1, MK-3207, BIBN-4096

## Abstract

Ewing sarcoma (EwS) is an aggressive cancer caused by chromosomal translocations generating fusions of the *EWSR1* gene with *ETS* transcription factors (in 85% *FLI1*). EWSR1-FLI1 induces gene expression via binding to enhancer-like GGAA-microsatellites, whose activity increases with the number of consecutive GGAA-repeats.

Herein, we investigate the role of the secretory neuropeptide CALCB (calcitonin related polypeptide β) in EwS, which signals via the CGRP-(calcitonin gene-related peptide) receptor complex, containing RAMP1 (receptor activity modifying protein 1) as crucial part for receptor specificity. Analysis of 2,678 gene expression microarrays comprising 50 tumor entities and 71 normal tissue types revealed that *CALCB* is specifically and highly overexpressed in EwS. Time-course knockdown experiments showed that *CALCB* expression is tightly linked to that of *EWSR1-FLI1*. Consistently, gene set enrichment analyses of genes whose expression in primary EwS is correlated to that of *CALCB* indicated that it is co-expressed with other EWSR1-FLI1 target genes and associated with signatures involved in stemness and proliferation. Chromatin immunoprecipitation followed by sequencing (ChIP-seq) data for EWSR1-FLI1 and histone marks from EwS cells demonstrated that EWSR1-FLI1 binds to a GGAA-microsatellite close to *CALCB*, which exhibits characteristics of an active enhancer. Reporter assays confirmed the strong EWSR1-FLI1- and length-dependent enhancer activity of this GGAA-microsatellite. Mass-spectrometry analyses of supernatants of EwS cell cultures demonstrated that CALCB is secreted by EwS cells. While short-term RNA interference-mediated *CALCB* knockdown had no effect on proliferation and clonogenic growth of EwS cells *in vitro*, its long-term knockdown decreased EwS growth *in vitro* and *in vivo*. Similarly, knockdown of *RAMP1* reduced clonogenic/spheroidal growth and tumorigenicity, and small-molecule inhibitors directed against the CGRP-receptor comprising RAMP1 reduced growth of EwS.

Collectively, our findings suggest that *CALCB* is a direct EWSR1-FLI1 target and that targeting the CALCB/RAMP1-axis may offer a new therapeutic strategy for inhibition of EwS growth.

## INTRODUCTION

Ewing sarcoma (EwS) is a malignant tumor of bone and soft tissue predominantly affecting children and adolescents^1^. Since specific treatment options do not exist, current therapy concepts comprise local surgery combined with conventional poly-chemotherapy and irradiation^1^. Despite such intense conventional therapy, prognosis of patients with metastatic disease still remains poor^2^. Thus, specific and, in particular, less toxic treatment options are urgently required.

EwS is characterized by gene fusions involving the *EWSR1* gene on chromosome 22 (chr22) and various members of the ETS family of transcription factors – most commonly *FLI1* on chr11 (85% of cases)^1^. *EWSR1-FLI1* can arise either through balanced chromosomal translocations or through complex genomic breakage/fusion events^3,4^. Notably, *EWSR1-FLI1* encodes an aberrant chimeric transcription factor, which binds DNA at ETS-like GGAA-motifs and furthermore at GGAA-microsatellites consisting of multiple sequential GGAA-motifs^5^. While EWSR1-FLI1 binding at single ETS-like motifs in gene promoters either activates or represses gene transcription, EWSR1-FLI1 binding at GGAA-microsatellites creates *de novo* enhancers, whose activity correlates positively with the number of consecutive GGAA-repeats^1,6,7^.

Although EwS is genetically well characterized, its precise cell of origin remains controversial. Transcriptome profiling and functional studies suggested that EwS may arise from mesoderm- or neural crest-derived mesenchymal stem cells^8,9^. Due to this histogenic uncertainty, there is currently no *bona fide* genetically engineered animal model available for EwS, which hampers the development of new therapeutic strategies^1,10^. In addition, recent sequencing efforts revealed, *EWSR1-ETS* translocations being virtually the only highly recurrent somatic mutation in EwS^11,12^. Like many other ligand-independent inducible transcription factor oncoproteins, also EWSR1-FLI1 proved to be notoriously difficult to target^1,13^. However, the EWSR1-FLI1-induced transcriptomic signature may harbor specific changes that could be exploited therapeutically.

To explore such EWSR1-FLI1 surrogate targets, we focused in this study on the putative EWSR1-FLI1 target gene *CALCB* (calcitonin related polypeptide β; alias CGRP2, calcitonin gene-related peptide 2), which encodes a neuropeptide that was already described in 1987 to be highly expressed in a EwS cell line^14,15^. Nevertheless, its functional effects in EwS have remained unexplored until now.

The *CALCB* gene is located next to its homolog *CALCA* (calcitonin related polypeptide α) on chr11p15.2 and encodes a secretory neuropeptide composed of 37 amino acids^16,17^. CALCB is predominantly expressed in the central nervous system and causes potent vasodilatation^18,19^. Signaling of both, CALCA and CALCB, is mediated through G protein coupled receptor complexes present on the cell surface. There is a variety of different receptors, formed by heterodimerization, which recognize both peptides. Most importantly they are recognized by the so called CGRP-receptor, which is formed by the calcitonin receptor-like receptor (CLR, encoded by the *CALCRL* gene) and RAMP1 (receptor activity-modifying protein 1). RAMP1 makes the receptor-complex specific for the binding of CALCA and CALCB ^20,21^. Receptor-ligand interaction leads to G protein mediated increase in intracellular cAMP-levels^22^. Apart from the above described CGRP-receptor, CALCB also binds to a receptor complex consisting of RAMP1 and the calcitonin receptor (CTR, encoded by the CALCR gene), which is called AMY1-(amylin subtype 1) receptor. However, this receptor is not specific for CALCA and CALCB but is also activated by binding of islet amyloid polypeptide (IAPP). Since the biological role of AMY1 is not fully understood, and given that both *CALCR* and *IAPP* are not or only barely expressed in EwS (**Supplementary Fig. S1**), we focused in this study on CALCB and the first CGRP-receptor containing CLR and RAMP1^21^.

Here, we show that *CALCB* is an EWSR1-FLI1 target gene highly overexpressed in EwS as compared to normal tissues and other childhood malignancies, and that its high expression is likely mediated through EWSR1-FLI1 binding to an enhancer-like GGAA-microsatellite. Proteomic and functional analyses revealed that CALCB, but not CALCA, is secreted by EwS cells and that suppression of either *CALCB* or its receptor *RAMP1* significantly reduced proliferation and clonogenic/spheroidal growth of EwS cells *in vitro*, as well as tumor growth *in vivo*, which can be mimicked *in vitro* by application of the small molecule CGRP-receptor inhibitors MK-3207 and BIBN-4096 (Olcegepant).

## MATERIALS & METHODS

### Analysis of microarray data

The microarray datasets for cancer and normal tissues were downloaded from public repositories and processed as described previously^23^. Data were generated on Affymetrix HG-U133Plus2.0 microarrays and normalized simultaneously by Robust Multi-chip Average (RMA) using brainarray chip description files (CDF; ENTREZg, v21) yielding one optimized probe-set per gene^24,25^. Accession codes of used datasets are given in **Supplementary Table 1**.

### Cell culture and provenience of cell lines

A673, HEK-293T, and SK-PN-DW cells were purchased from the American Type Culture Collection (ATCC, Manassas, VA, USA; CRL-1598, CRL-1573, and CRL-2139, respectively). RDES, SK-ES1, SK-N-MC, and MHH-ES1 cells were provided by the German collection of Microorganism and Cell lines (DSMZ, Braunschweig, Germany). TC-71 cells were kindly provided by the Children’s Oncology Group (COG, Monrovia, CA, USA) and ES7, EW-1, EW-3, EW-7, EW-16, EW-18, EW-22, EW-24, LAP35, MIC, ORS, POE, SKNPLI, and STA-ET1 cells were provided by O. Delattre (Institute Curie, Paris, France). SB-KMS-KS1 was established in the Department of Pediatrics at the TU Munich (Munich, Germany) and described previously^26^. A673/TR/shEF1 cells, which contain a doxycycline (dox)-inducible shRNA against EWSR1-FLI1, were kindly provided by J. Alonso (Madrid, Spain)^27^. All cell lines were grown at 37°C and 5% CO_2_ in a humidified atmosphere. RPMI 1640 medium with stable glutamine (Biochrom, Berlin, Germany), 10% tetracycline-free FCS (Biochrom), 100 U/ml penicillin (Biochrom), and 100 μg/ml streptomycin (Biochrom) was used as standard medium to grow the cells. For cell lines, which tend to grow in suspension, TPP cell culture flasks (Faust, Klettgau, Germany) were coated with 1:40 PBS (Biochrom) diluted collagen solution (Sigma-Aldrich/Merck Millipore, Darmstadt, Germany) in order to keep them attached to the flasks. Cells were routinely checked for mycoplasma infection by nested PCR. Cell line purity was confirmed by STR profiling.

### RNA extraction, reverse transcription, and quantitative real-time PCR (qRT-PCR)

RNA for analysis of gene expression with qRT-PCR from cell lysates and frozen tumor tissue was extracted using the NucleoSpin RNA Kit (Macherey-Nagel, Düren, Germany). Subsequent reverse transcription was performed with the High Capacity cDNA Reverse Transcription Kit (Thermo Fisher Scientific, Waltham, MA, USA) utilizing 1 μg RNA per reaction and following manufacturers’ protocol of both kits. qRT-PCRs were performed using SYBR green (Applied Biosystems, Waltham, MA, USA) with a total volume of 15 μl. cDNA was diluted 1:10 and concentration of primers was 0.5 μM. Oligonucleotides were purchased from MWG Eurofins Genomics (Ebersberg, Germany). Expression levels were determined with the CFX Connect Real time PCR Cycler (Bio-Rad Laboratories, Hercules, CA, USA) in a two-step protocol: initial enzyme activation and denaturation at 95°C for 2 min, denaturation at 95°C for 10 sec, and annealing and extension at 60°C for 20 sec (repeating the last two steps 49 times), followed by a melting curve starting at 55°C and increasing by 0.5°C every 10 sec until a temperature of 95°C was reached. Expression levels were calculated according to the 2^−ΔΔCT^ method^28^. *RPLP0* served as housekeeping gene. Primer sequences were as follows:

*RPLP0* forward, 5’-GAAACTCTGCATTCTCGCTTC-3’;

*RPLP0* reverse, 5’-GGTGTAATCCGTCTCCACAG-3’;

*EWSR1-FLI1* forward, 5’-GCCAAGCTCCAAGTCAATATAGC-3’;

*EWSR1-FLI1* reverse, 5’-GAGGCCAGAATTCATGTTATTGC-3’;

*CALCB* forward, 5’-GCTCTCAGTATCTTGGTCCTG-3’;

*CALCB* reverse, 5’-CACATAGTCCTGCACCAGTG-3’;

*RAMP1* forward, 5’-CCCAGTTCCAGGTAGACATG-3’;

*RAMP1* reverse, 5’-CCAGCTTCTCCGCCATGTG-3’.

### Quantification of *CALCB* gene expression levels in EwS cell lines

Different EwS cell lines were cultured under standard conditions. After a minimum of 48 h of culture, cells were harvested at a confluency of approximately 80%, and RNA extraction was carried out. *CALCB* expression was determined using qRT-PCR as described. *CALCB* expression levels were calculated relative to that of the A673 EwS cell line.

### Quantification of EWSR1-FLI1-dependent *CALCB* gene expression *in vivo*

For analysis of *in vivo CALCB* expression depending on EWSR1-FLI1, 5 × 10^6^ A673/TR/shEF1 EwS cells, which harbor a dox-inducible shRNA against *EWSR1-FLI1*, were injected subcutaneously in the flanks of immunocompromised NSG (Nod/scid/gamma) mice. When tumors reached an average volume of 180 mm^3^, mice were randomized and either received 2 mg/ml dox (Beladox, bela-pharm, Vechta, Germany) and 5% sucrose (Sigma-Aldrich/Merck Millipore) in the drinking water (dox +) or only 5% sucrose (dox −). Mice were sacrificed 96 h after beginning of dox-treatment, and tumors were collected for RNA analysis. Total RNA was extracted using the ReliaPrep miRNA Cell and Tissue Miniprep System (Promega, Madison, WI, USA). Knockdown of *EWSR1-FLI1* was confirmed by qRT-PCR and proved *EWSR1-FLI1* expression to be downregulated onto 15% of the control (dox −). Tumor purity (> 95%) was confirmed in routine histology (hematoxylin and eosin [H&E] stains). The transcriptomes of 3 tumors of each group were profiled on Affymetrix Clariom D arrays (RIN > 9). Microarray data were simultaneously normalized on gene level using Signal Space Transformation Robust Multi-Chip Average (SST-RMA) and Affymetrix CDF as described^29^.

### Analysis of chromatin-immunoprecipitation followed by sequencing (ChIP-Seq) data

Publicly available data were retrieved from the Gene Expression Omnibus (GEO; GSE61944)^7^ and the ENCODE project^30^, processed as described^31^, and displayed in the UCSC genome browser. The used samples are available under the following accession codes:

GSM736570 ENCODE_SKNMC_hg19_DNAseHS_rep2;

GSM1517546 SKNMC.shGFP96.FLI1;

GSM1517555 SKNMC.shFLI196.FLI1;

GSM1517548 SK-N-MC_shGFP_96h_H3K4me1;

GSM1517557 SK-N-MC_shFLI1_96h_H3K4me1;

GSM1517547 SKNMC.shGFP96.H3K27ac;

GSM1517556 SKNMC.shFLI196.H3K27ac;

GSM1517569 A673.shGFP48.FLI1;

GSM1517572 A673.shFLI148.FLI1;

GSM1517571 A673.shGFP96.H3.k27ac;

GSM1517574 A673.shFLI196.H3K27ac.

### Dual luciferase reporter assays

A 359 base pairs (bp) fragment around the *CALCB*-associated GGAA-microsatellite was cloned from two different EwS cell lines (TC-71 and MHH-ES1) by PCR into the pGL3 Luciferase Reporter Vector (Promega) using the In-Fusion HD Cloning Kit (Clontech, Takara Bio USA, CA, USA). The DNA from the cell lines was extracted using the NucleoSpin Tissue genomic DNA prep kit (Macherey-Nagel) and digested with the restriction enzymes Eco-RV and SphI (New England Biolabs [NEB], Ipswich, MA, USA), followed by a fragment separation by gel electrophoresis, and fragment purification of the band at 8,200 bp using the NucleoSpin Gel and PCR Clean-up kit (Macherey-Nagel). A touch-down PCR was performed with 100 ng of the purified DNA fragment using the Infusion-Primers:

forward: 5’-ctagcccgggctcgagGAGCCCTTTAGTATCCCCTTTG-3’;

reverse: 5’-gatcgcagatctcgagACCCTTGT ACTAACAT GCTTCG-3’

(MWG Eurofins Genomics) at a concentration of 0.5 μM, and the components of the Go Taq Hot Start Polymerase (Promega) according to the manufacturer’s instructions. The thermal protocol was as follows: activation at 95°C for 2 min, followed by 20 cycles of 98°C for 10 sec, 59-49°C for 30 sec (decreasing 0.5°C in each cycle), and 72°C for 1 min, finalizing at 72°C for 5 min. The PCR product was separated by gel electrophoresis, and the desired fragment of 359 bp was purified with the NucleoSpin Gel and PCR Clean-up kit (Macherey-Nagel). The infusion reaction with the In-Fusion HD Cloning Kit (Clontech) was initiated using 10 ng of the linearized pGL3 Luciferase Reporter Vector (Promega), digested with XhoI (NEB) and prepared according to the manual, and 20 ng of the DNA fragment. The reaction was incubated at 50°C for 15 min, stopped on ice for 5 min, and transformed into *E. coli* Stellar competent cells (Clontech) using the DNA of the infusion mixture. Bacteria were grown on agar plates supplemented with ampicillin at a final concentration of 100 μg/ml (Sigma-Aldrich/Merck Millipore) overnight at 37°C. Colonies were picked on the next morning and checked for clones containing the plasmid with the insert by colony PCR. Positive clones were incubated overnight in LB broth (Miller)-medium with 100 μg/ml ampicillin (Sigma-Aldrich/Merck Millipore) at 37°C, and plasmids were purified using the PureYield Plasmid Midiprep System 2 (Promega). The identity of the fragment was validated by Sanger sequencing (Eurofins GATC Biotech, Konstanz, Germany).

For the luciferase reporter assay, 2 × 10^5^ A673/TR/shEF1 EwS cells, harboring a dox-inducible shRNA against *EWSR1-FLI1*, were plated out in a well of a 6-well plate (TPP, Faust) in 1.8 ml of growth medium and transfected with the microsatellite-containing pGL3-luc vector and *Renilla* pGL3-Rluc vector (ratio, 100:1) using Lipofectamine LTX with Plus Reagent (Thermo Fisher Scientific). Transfection media was replaced by media with/without dox (1 μg/ml; VWR/Merck, Radnor, PA, USA) 4 h after transfection. After 72 h, the cells were lysed and assayed with a dual luciferase assay system (Berthold Technologies, Bad Wildbad, Germany). Firefly luciferase activity was normalized to Renilla luciferase activity.

### Gene set enrichment analysis (GSEA)

To identify gene signatures and biological processes associated with *CALCB* in normalized gene expression data from 166 primary EwS^32^, GSEA was performed on lists of genes ranked by their correlation coefficient with *CALCB* (MSigDB, c2.cpg.v6.2). GSEA was carried out with 1,000 permutations in default settings^33^.

### Generation of cell lines with dox-inducible shRNAs

For generation of dox-inducible EwS cell lines (here in A673 and RDES), either a non-targeting negative control shRNA (MWG Eurofins Genomics) or specific shRNAs targeting *CALCB* or *RAMP1* (MWG Eurofins Genomics) were cloned in the pLKO-Tet-on-all-in-one system (Addgene plasmid # 21915, Cambridge, MA, USA) as described previously^34^. Lentivirus production was performed in HEK-293T cells. A673 and RDES EwS cells were infected with the respective lentiviruses and selected with 1.5 μg/ml puromycin (Invivogen, Toulouse, France). Single cell cloning was performed, and knockdown efficacy of individual clones was assessed by qRT-PCR 48 h after addition of dox (1 μg/ml; VWR/Merck). The shRNA target sequences were as follows:

shControl, 5’-CAACAAGATGAAGAGCACCAA-3’;

shCALCB1, 5’-AAGGAAT GAAACTGAAT GCAA-3’;

shCALCB4, 5’-AACCTTGGTGATGCATTACAA-3’;

shRAMP1_3, 5’-GCGCACTGAGGGCATTGTGTA-3’;

shRAMP1_4, 5’-TGCCTGCCAGGAGGCTAACTA-3’.

### Proliferation assays

1-5 × 10^5^ cells were seeded in 6-well plates (TPP, Faust) in 1.5 ml of standard growth medium. Gene knockdown was induced by addition of 1 μg/ml dox (VWR/Merck) to the growth medium (refreshed every 48-72 h) to cells harboring an inducible shRNA against *CALCB* or by serial transfections with 25 nM siRNA directed against *CALCB* (Hs_CALCB_1 FlexiTube siRNA or Hs_CALCB_4 FlexiTube siRNA, QIAGEN, Hilden, Germany) or a scrambled control siRNA (MISSION siRNA Universal Negative Control #1, Sigma-Aldrich/ Merck Millipore) following the manufactures handbook of the transfection reagent HiPerfect (QIAGEN). After 2-4 h, 3 ml of standard growth medium was added to prevent toxic effects of the transfection reagent. After 24 h, the medium was exchanged and after another 24 h a second transfection was performed. Cell counts were determined by using standardized hemocytometers (C-chips, Biochrom) and Trypan-blue (Sigma-Aldrich/Merck Millipore) exclusion 72 h after seeding in the short-term proliferation assay and 6-9 days after seeding in the long-term proliferation assay.

### Colony-forming assay

A673 and RDES EwS cells harboring a dox-inducible shRNA construct against *CALCB* or *RAMP1* or RDES wildtype cells were seeded at concentrations of 100-1,000 cells per well of 12-well plates and grown in standard culture medium for 12-14 days. Cells were treated with/without 1 μg/ml dox (VWR/Merck) and RDES wildtype cells were serially transfected with a non-targeting siRNA or siRNAs against *CALCB* (as described above). Colonies were stained with crystal violet (Sigma-Aldrich/Merck Millipore) and the number of colonies was quantified using ImageJ.

### Sphere-formation assay

For analysis of 3D sphere formation, A673 and RDES EwS cells harboring a dox-inducible shRNA construct against *CALCB* or *RAMP1* were seeded at a density of 1,000 cells/well of ultra-low attachment 96-well plates (Corning, NY, USA) in 80 μl standard cell culture medium with/without dox (1 μg/ml; VWR/Merck). The culture medium was refreshed by adding 10 μl medium with/without dox on top every second day. Spheroidal growth was monitored for 14 days. Thereafter, phase-contrast imaging and morphological analyses of spheres were carried out with an inverted Zeiss Axiovert 25 microscope (Jena, Germany) equipped with a Zeiss Axiocam 105 color camera (Aptina CMOS Color Sensor, square pixels of 2.2 μm side length, 2,560 × 1,920 pixel resolution). Sphere numbers and diameters were analyzed with ImageJ.

### Analysis of tumor growth *in vivo*

2.5 × 10^6^ A673 EwS cells harboring a dox-inducible shRNA construct against *CALCB* or *RAMP1* or a non-targeting control shRNA (shControl) were injected subcutaneously in NSG mice. After 10-14 days, when tumors were first palpable, mice were randomized and thereafter received either 2 mg/ml dox (bela-pharm) dissolved in 5% sucrose (Sigma-Aldrich/Merck Millipore) and sterile water (dox +) or 5% sucrose in sterile water alone (dox −). Tumor growth was monitored with a caliper every other day and mice were sacrificed by cervical dislocation when the tumors exceeded an average diameter of 15 mm (prior start of the experiment defined as “event”). Experiments were approved by local authorities and conducted in accordance with the recommendations of the European Community (86/609/EEC) and UKCCCR (guidelines for the welfare and use of animals in cancer research).

### Small-molecule inhibitor assays

A673 and A673/TR/shRAMP1_4 EwS cells, latter harboring a dox-inducible shRNA against *RAMP1*, were seeded at a density of 1,500 cells per well of a 96-well plate (TPP, Faust) in 50 μl standard growth medium with/without dox (1 μg/ml; VWR/Merck). After 24 h of incubation, treatment was started by addition of 50 μl standard growth medium containing either different concentrations of the CGRP-receptor inhibitor MK-3207 (AdooQ Bioscience, Irvine, CA, USA) dissolved in DMSO (Sigma-Aldrich/Merck Millipore) or the corresponding concentration of DMSO without inhibitor and dox refreshment for dox + wells. After 72 h, read-out was performed by addition of 20 μl of 1:10 dissolved Resazurin (Sigma-Aldrich/Merck Millipore) to the cells and measurement of cell activity with a plate reader (Thermo Fisher Scientific) after 7 h of incubation. For analysis of 2D colony-formation capacity under inhibitor treatment, A673 and RDES EwS cells were seeded at a density of 100 cells/well of 12-well plates (TPP, Faust) in a volume of 1 ml culture medium. 48 h after seeding, inhibitors were added at final concentrations of 100 μM for BIBN-4096 (Olcegepant; R&D systems, Minneapolis, MN, USA) and 20 μM for MK-3207 (AdooQ Bioscience). DMSO (Sigma-Aldrich/Merck Millipore) served as a control. After 1-3 weeks, colonies were gently washed twice with PBS (Biochrom) and stained with 500 μl crystal violet solution (Sigma-Aldrich/Merck Millipore). Colonies were photographed, and the number of colonies was counted using Image J.

For analysis of 3D sphere-formation capacity under inhibitor treatment, A673 and RDES EwS cells were seeded at a density of 1,000 cells/well in 80 μl culture medium in wells of 96-well ultra-low attachment culture plates (Corning). After 24 h of incubation, 20 μl culture medium containing either inhibitor or DSMO (control; Sigma-Aldrich/Merck Millipore) was added to the wells resulting in a final concentration of 100 μM BIBN-4096 (Olcegepant; R&D systems) or 20 μM MK-3207 (AdooQ Bioscience), respectively. After 14 days, spheres were photographed, and their number and size were analyzed using ImageJ.

### Human samples and immunohistochemistry (IHC)

Available tissue microarrays (TMA) of primary EwS tumors containing 2 cores of each sample, with a diameter of 1 mm, as well as internal controls were stained for CALCB. Analysis were carried out in approval with the LMU Munich ethics committee.

For IHC, 4-μm sections were cut and antigen retrieval was carried out by heat treatment using target unmasking fluid (PanPath, Budel, Netherlands). Slides were incubated for 60 min at room temperature with a rabbit polyclonal anti-CALCB antibody (bs-0791R, Bioss Antibodies Inc., MA, USA; dilution 1:120). Then slides were incubated with a secondary anti-rabbit IgG antibody (Vectastain ABC-Kit Elite Universal, Vector laboratories, Burlingame, CA, USA) followed by target detection using DAB plus (Agilent Technologies, Santa Clara, CA, USA). Counterstaining was performed with Hematoxylin Gill’s Formula (Vector). Intensity of CALCB staining was scored independently by two researchers in a scale from 0 to 2 (0 = majority of the cells is negative for CALCB staining, 1 = majority of the cells shows moderate CALCB staining, and 2 = majority of the cells shows strong CALCB staining).

### CD31 staining and evaluation of microvessel density

For CD31 staining, 4 μm sections of formalin-fixed and paraffin-embedded tumor tissue derived from EwS xenografts in mice were cut, and heat treated using the Target Retrieval Solution (Agilent Technologies). Thereafter, tissue slides were stained with a primary monoclonal rat anti-CD31-antibody (DIA-310, Dianova, Hamburg, Germany; dilution 1:150, 60 min incubation at room temperature). A secondary biotinylated and mouse-absorbed anti-rat-IgG-antibody (BA 4001, Vector; dilution 1:100) was used. After Streptavidin HRP (Novocastra Laboratories, Newcastle upon Tyne, United Kingdom) treatment, DAB plus (Agilent Technologies) was used for target detection. The slides were counterstained with Hematoxylin Gill’s Formula (Vector). For evaluation of the microvessel density in the CD31 stained slides, the Chalkley-grid method was used^35^. To this end, the number of overlaps of a CD31 positive cell with a dot of the Chalkley-grid in each quarter of the grid in 4 independent regions of the CD31 stained slide was counted, and the mean vessel density of the tumor was calculated.

### Mass spectrometry analyses

A673 EwS cells were seeded at a density of 4 × 10^6^ cells per T150 flask (TPP, Faust) in 20 ml of standard culture medium. After 48 h, the supernatant from the cells was removed, and the cells were washed with PBS (Biochrom) twice. Thereafter, the cells were grown for further 24 h in 20 ml Opti-MEM (Thermo Fisher Scientific) only. Afterwards, the supernatants were collected and immediately frozen at −80°C until the mass spectrometry was performed. For mass spectrometry analysis, samples were lyophilized and resuspended in 50 mM ammonium bicarbonate. 200 μg of protein were reduced and alkylated using 10 mM DTT (Thermo Fisher Scientific) and 100 mM iodoacetamide (Sigma-Aldrich/Merck Millipore), respectively. Next, samples were digested using 20 ng/μl trypsin (NEB) for 18 h at 37°C. Samples were separated in a Nano-HPLC (NanoLC-2D, Eksigent, Sciex, CA, USA) using a C18 column and a gradient composed of solvent A (5% acetonitrile) and solvent B (95% acetonitrile). The program was: 5% acetonitrile for 5 min, 5-100% for 50 min, and 100% for 10 min. Eluted samples were spotted (Eksigent) on a 384-well plate and 1 μl of the matrix α-cyano-4-hydroxycinnamic acid (10 mg/ml in 50% acetonitrile) was added. The mass spectrometry analysis was performed on a MALDI-TOF/TOF 4800 (Sciex) using positive mode. The data were analyzed with the Trans-Proteomic Pipeline (Seattle Proteome Center, WA, USA).

## RESULTS

### *CALCB* is an EWSR1-FLI1 target gene highly but heterogeneously expressed in EwS

In the search of potential EWSR1-FLI1 surrogate targets, we analyzed publicly available gene expression microarray data comprising 71 normal tissue types and 50 tumor entities, which identified *CALCB* as being highly overexpressed in EwS compared to most tumor entities and all normal tissues except for tissue derived from trigeminal ganglia (**Figure 1A, B; Supplementary Fig. S2**). The high expression of CALCB in EwS was validated by immunohistochemical staining of a TMA of primary EwS tumors, of which 44% (39/89) displayed a high and 37% (33/89) an intermediate immunoreactivity score (2 or 1, respectively) for CALCB expression (**Supplementary Fig. S3**).

**Figure 1:**
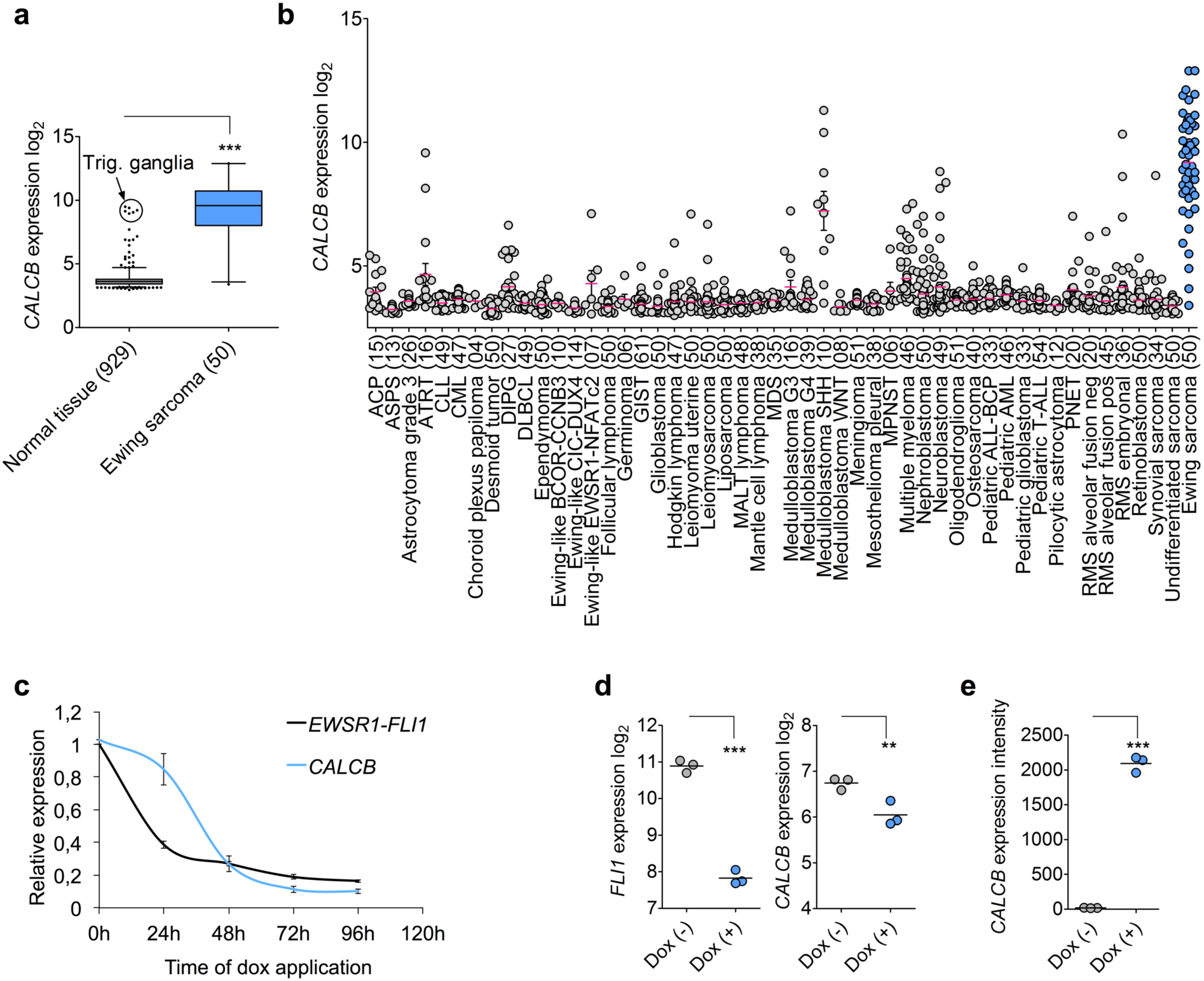
*CALCB* is an EWSR1-FLI1 target gene specifically expressed in EwS. **A)** Analysis of *CALCB* expression in EwS (*n* = 50) and normal tissues (71 tissue types, *n* = 929 samples). Data are represented as box-plots. Horizontal bars indicate median expression levels, boxes the interquartile range. Whiskers indicate the 2.5^th^ and 97.5^th^ percentile, respectively. Unpaired two-tailed student’s t-test. **B)** Analysis of *CALCB* expression in EwS (*n* = 50) and different (pediatric) tumors (49 types, *n* = 1 699 samples). Publicly available microarray data are represented as dot plots in log2-scale with mean and SEM. Each dot represents one sample. The number of samples is given in parentheses. EwS highlighted in blue color. **C)** Time-course analysis of *CALCB* and *EWSR1-FLI1* expression in A673/TR/shEF1 EwS cell harboring a dox-inducible shRNA against EWSR1-FLI1 by qRT-PCR *in vitro* after dox application. Given are mean expression levels and SEM (*n* = 7). **D)** Analysis of *CALCB* expression in xenografts derived from A673/TR/shEF1 cells with/without dox-treatment for 96 h *in vivo*. Gene expression levels were determined by Affymetrix Clariom D microarrays as previously described^29^. Expression levels are shown in log_2_-scale; horizontal bars indicate mean expression level (*n* = 3); unpaired two-tailed student’s t-test. **E)** Analysis of *CALCB* expression in a published dataset (GSE64686)^38^ with ectopic *EWSR1-FLI1* expression in human embryoid bodies. Data were generated on Affymetrix HG-U133Plus2.0 microarrays and normalized simultaneously by RMA and brainarray CDF (ENTREZg; v21). Horizontal bars indicated mean expression levels; unpaired two-tailed student’s t-test. n.s. *P* > 0.05; * *P ≤* 0.05; ** *P ≤* 0.01; *** *P ≤* 0.001

This EwS-specific expression pattern suggested a potential regulatory relationship between EwS-specific *EWSR1-ETS* fusion oncogenes and *CALCB*. To test this hypothesis, we performed time-course knockdown experiments of *EWSR1-FLI1* in the EwS cell line A673/TR/shEF1, which harbors a dox-inducible shRNA against *EWSR1-FLI1* and measured the expression of *EWSR1-FLI1* and *CALCB* at different time points after start of dox-treatment (0-96 h). The results showed that the expression of *CALCB* is tightly linked to that of *EWSR1-FLI1* (**Fig. 1C**), which was confirmed in xenografts derived from A673/TR/shEF1 cells *in vivo* (**Fig. 1D**). Conversely, ectopic expression of *EWSR1-FLI1* in human embryoid bodies strongly induced *CALCB* expression (**Fig. 1E**).

To further assess this regulatory relationship, we explored available EWSR1-FLI1 ChIP-seq data from two EwS cell lines (A673 and SK-N-MC) and found strong EWSR1-FLI1 binding at intron 5 of the longest isoform (isoform 3) of the *CALCB* gene, which mapped to a GGAA-microsatellite that showed epigenetic characteristics of an active enhancer (**Fig. 2A**). Knockdown of EWSR1-FLI1 in both cell lines abolished the EWSR1-FLI1 signal at this GGAA-microsatellite and markedly reduced the signals for acetylated H3K27 (H3K27ac), indicating abrogated enhancer activity upon EWSR1-FLI1 silencing (**Fig. 2A**).

**Figure 2:**
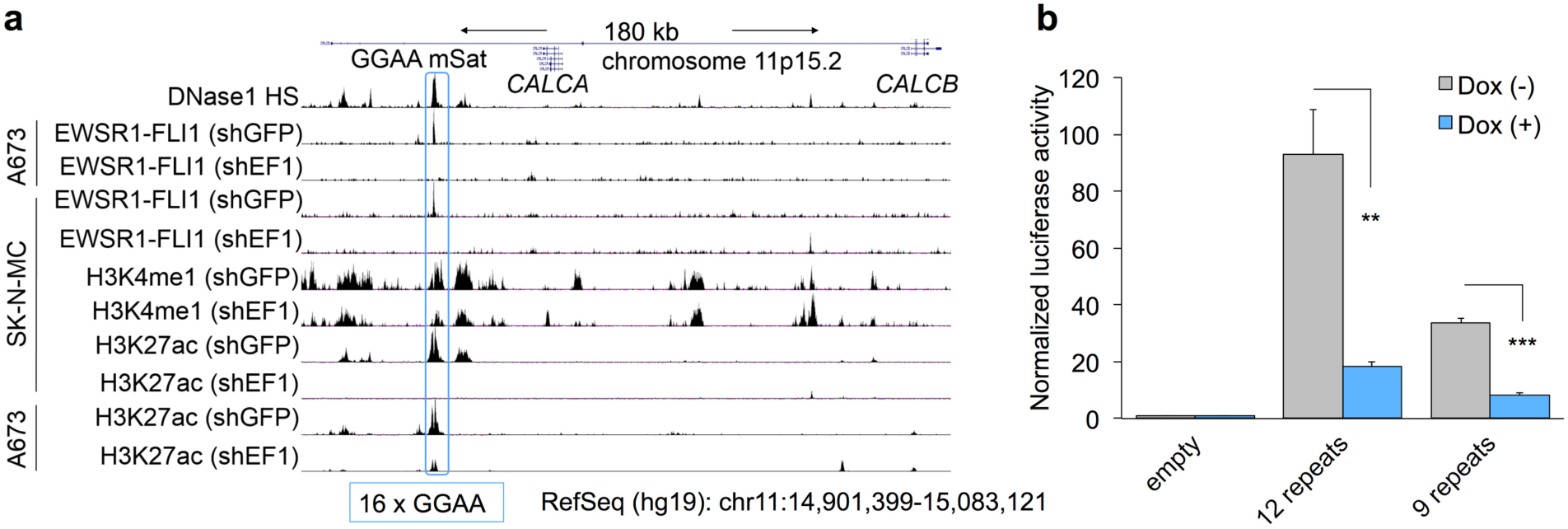
*CALCB* expression is regulated through EWSR1-FLI1 binding to a nearby enhancer-like GGAA-microsatellite. **A)** Integrative genomic view of published ChIP-seq^7^ and DNAse-seq data^30^ of the *CALCB* locus. Data were generated in A673 and SK-N-MC EwS cells, stably transfected with either shRNA targeting *GFP* (shGFP; negative control) or *EWSR1-FLI1* (shEF1). The blue box marks the location of the *CALCB* gene associated GGAA-microsatellite, consisting of 16 repeats of the GGAA-motif in the human reference genome (hg19). **B)** Luciferase reporter assays in A673/TR/shEF1 cells with/without knockdown of *EWSR1-FLI1* (Dox +/−) 72 h after transfection with plasmids containing a 359 bp fragment around the CALCB-associated GGAA-microsatellite as displayed in **Fig. 2A**. Data are presented as mean (*n* = 3-5) and SEM; unpaired two-tailed student’s t-test. n.s. *P* > 0.05; * *P ≤* 0.05; ** *P ≤* 0.01; *** *P ≤* 0.001

To confirm its EWSR1-FLI1-dependent enhancer activity, we cloned a 359 bp fragment containing this GGAA-microsatellite from two EwS cell lines (TC-71 and MHH-ES1) into the pGL3 luciferase reporter vector and performed reporter assays in A673/TR/shEF1 cells with/without silencing of *EWSR1-FLI1*. In these assays, we observed strong enhancer activity of the GGAA-microsatellite, which was significantly diminished upon *EWSR1-FLI1* knockdown (**Fig. 2B**). In accordance with the higher number of consecutive GGAA-repeats and higher *CALCB* expression levels in TC-71 EwS cells (12 repeats) as compared to MHH-ES1 EwS cells (9 repeats), we noted a higher enhancer activity of the GGAA-microsatellite derived from TC-71 as compared to the microsatellite derived from MHH-ES1 in luciferase assays (**Fig. 2B, Supplementary Fig. S4**). Collectively, these data provide evidence that *CALCB* is a direct EWSR1-FLI1 target gene, whose high but heterogeneous expression in EwS is regulated by EWSR1-FLI1 binding to an intronic, polymorphic, and enhancer-like GGAA-microsatellite.

### *CALCB* expression in primary EwS correlates with proliferation signatures

To obtain first clues on the potential functional role of CALCB in EwS, we performed gene set enrichment analysis (GSEA) on *CALCB* co-expressed genes in a transcriptome dataset of 166 primary EwS^32^. GSEA revealed that *CALCB* is co-expressed with other EWSR1-FLI1 target genes (ZHANG_TARGETS_OF_EWSR1-FLI1_FUSION)^36^ and with gene signatures involved in stemness and proliferation (**Fig. 3A, B**).

**Figure 3:**
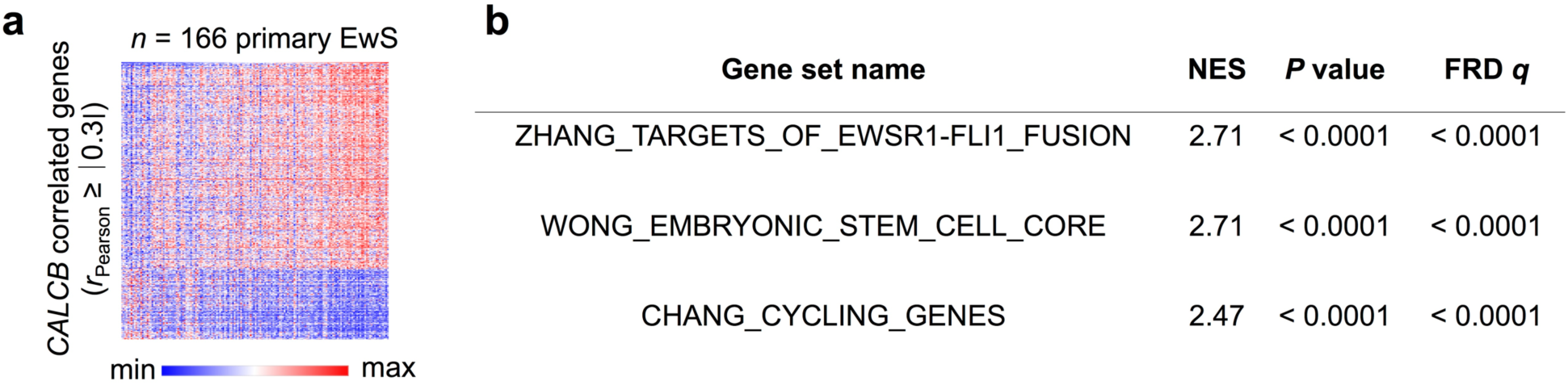
*CALCB* expression in primary EwS correlates with proliferation signatures. **A)** Heatmap of CALCB correlated genes (*r*_Pearson_ ≥ |0.3|) in 166 primary EwS tumors. **B)** Results of the gene set enrichment analysis (GSEA) on the ranked list of *CALCB* correlated genes as in (A). NES, normalized enrichment score.

### CALCB signaling in EwS cells contributes to growth of EwS

To test the bioinformatic predictions from our GSEA in primary EwS, we carried out several functional experiments in EwS models. Mass spectrometry analysis showed that CALCB was readily detectable in FCS-free cell culture supernatants conditioned by A673 EwS cells, whereas it was not detectable in FCS-free cell culture medium not conditioned by EwS cells (**Supplementary Table 2**), suggesting that CALCB is indeed secreted by EwS cells. Notably, in accordance with the low expression levels of *CALCA* in primary EwS cells (**Supplementary Fig. S1**), CALCA was not detectable in cell culture supernatants of EwS cells (**Supplementary Table 2**).

To investigate the functional role of CALCB in EwS, we performed RNA interference experiments in two EwS cell lines (RDES and A673), which showed relatively high or moderate *CALCB* expression levels as compared to 20 other EwS cell lines (**Supplementary Fig. S4**). While the short-term knockdown of *CALCB* for 3 days had no effect on cellular proliferation (**Supplementary Fig. S5**), its long-term knockdown for 6-9 days significantly reduced proliferation and clonogenic growth *in vitro* (**Fig. 4A, B**). As no differences in the relative number of dead cells was detectable by Trypan-blue labeled cell counting, the reduced proliferation appeared to be not mediated by an increase in cell death (**Fig. 4A**).

**Figure 4:**
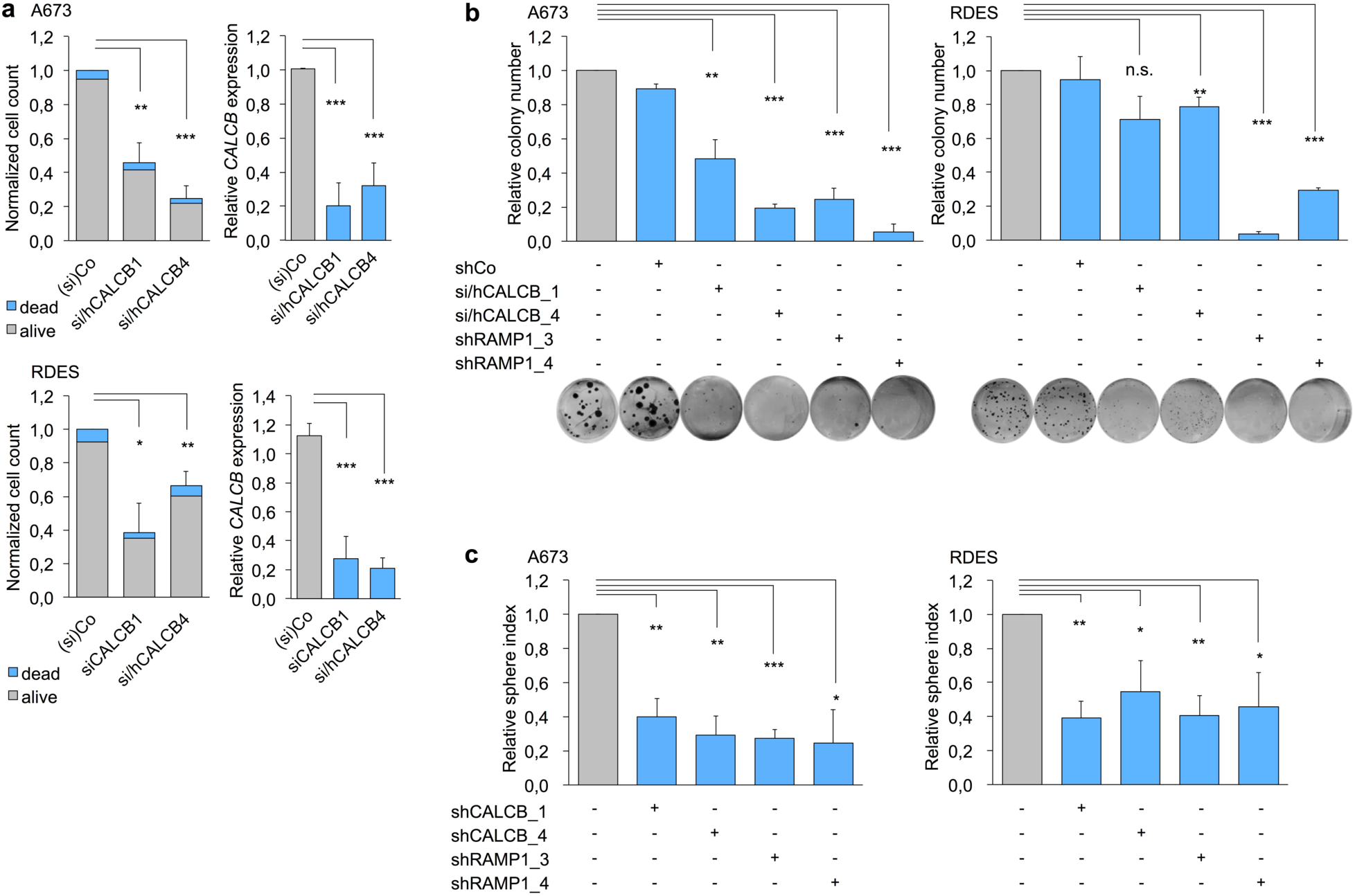
Knockdown of *CALCB* or *RAMP1* inhibits proliferation of EwS cells *in vitro*. **A)** Analysis of cell proliferation and cell death of A673 and RDES EwS cells with/without siRNA- or shRNA-mediated knockdown of *CALCB*. Left panel A673 and RDES: Given is the mean of relative cell count compared to Co (Control) either received according doses of a non-targeting siControl or did not receive dox in assays with dox-inducible shRNAs (*n* = 3-5). SEM and unpaired two-tailed student’s t-test of relative total cell count. Right panel A673 and RDES: Knockdown of *CALCB* was verified by qRT-PCR. Given is the mean gene expression and SEM; unpaired two-tailed student’s t-test. **B)** Colony-forming assays of A673 and RDES EwS cells with/without siRNA- or dox-inducible shRNA-mediated knockdown of *CALCB* or *RAMP1*. Mean colony number and SEM normalized to control, which either received according doses of a non-targeting siControl or did not receive dox in assays with dox-inducible shRNAs (*n* = 3-6). Unpaired two-tailed student’s t-test. Representative images of each condition are shown. **C)** Sphere formation assays of A673 and RDES EwS cells with/without dox-inducible shRNA-mediated knockdown of *CALCB* and *RAMP1*. Sphere index was calculated by addition of diameter of all existing spheres in one well divided by diameter of spheres in the control well. Mean and SEM (*n* = 3-6); unpaired two-tailed student’s t-test. n.s. *P* > 0.05; * *P ≤* 0.05; ** *P ≤* 0.01; *** *P ≤* 0.001

Interestingly, knockdown of *RAMP1* – the crucial component of the CALCB receptor complex – phenocopied the effect of *CALCB* knockdown in clonogenic growth assays (**Fig. 4B**) as well as in 3D sphere formation assays (**Fig. 4C**).

We next investigated whether knockdown of either gene could alter growth of xenografted EwS cells *in vivo*. To this end, we injected A673 cells, harboring a dox-inducible shRNA against either *CALCB* or *RAMP1*, subcutaneously in NSG mice. When tumors were palpable, we induced the knockdown of the respective gene by addition of dox to the drinking water. In both settings, knockdown of the corresponding gene significantly delayed tumor growth, which led to a delayed achievement of a mean tumor diameter of 15 mm, that was defined as termination criteria before, and therefore allowed a prolonged survival of the animals (**Fig. 5A, B**). However, dox-treatment of mice carrying tumors with dox-inducible expression of a non-targeting control shRNA did not alter tumor growth compared to mice not receiving dox (data not shown), as also described previously for other xenografted EwS cell lines^31^. Although *CALCB* and *RAMP1* were knocked down to low levels as confirmed by qRT-PCR of tumor tissue (**Fig. 5A, B**), the growth-inhibiting effect was more pronounced in the group of the *RAMP1* knockdown.

**Figure 5:**
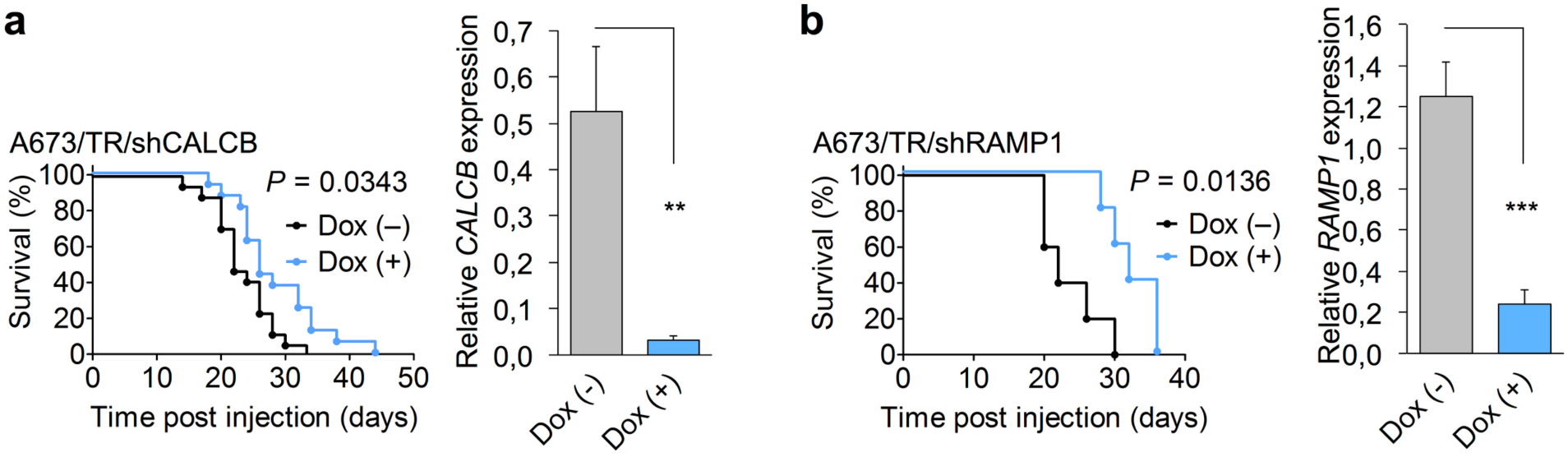
Knockdown of *CALCB* or *RAMP1* inhibits proliferation of EwS cells *in vivo*. **A)** Left panel: Analysis of tumor growth of A673 EwS cells with/without dox-inducible knockdown of *CALCB* in NSG mice (*n* = 33). Event was defined as average diameter of 15 mm. Event-free survival time of mice was analyzed by the Kaplan-Meier-method and a log rank test. Right panel: Knockdown of *CALCB* in the tumors of dox treated mice was verified by qRT-PCR. Given are mean normalized gene expression levels and SEM; unpaired two-tailed student’s t-test. **B)** Left panel: Analysis of tumor growth of A673 EwS cells with/without dox-inducible knockdown of *RAMP1* in NSG mice (*n* = 10). Event was defined as average diameter of 15 mm. Event-free survival time of mice was analyzed by the Kaplan-Meier-method and a log rank test. Right panel: Knockdown of *RAMP1* in the tumors of dox treated mice was verified by qRT-PCR. Given are mean normalized gene expression levels and SEM; unpaired two-tailed student’s t-test. n.s. *P* > 0.05; * *P ≤* 0.05; ** *P ≤* 0.01; *** *P ≤* 0.001

Taken together, these data suggest that CALCB is a secreted peptide in EwS and that the CALCB/RAMP1-axis promotes growth of EwS cells.

### Pharmacological inhibition of the CALCB/RAMP1-axis decreases growth of EwS cells

To test whether the CALCB/RAMP1-axis could also be exploited therapeutically in EwS, we treated EwS cells with the small molecule CGRP-receptor inhibitor MK-3207 for 3 days and quantified cell viability with a Resazurin-assay. For these assays, we used dox-inducible *CALCB* or *RAMP1* knockdown EwS cells and applied increasing doses of MK-3207. We observed a dose-dependent reduction of cell viability (**Fig. 6A**), which could be partially abrogated by knockdown of *RAMP1* – the central component of the inhibitor’s target structure (**Fig. 6B**). These data suggest that albeit relatively high doses of MK-3207 were applied to reduce viability of EwS cells, its effect was specific for the CALCB/RAMP1-axis. To validate these findings, we performed colony- and sphere-formation assays under MK-3207 treatment and replicated these experiments with another, well known small molecule CGRP inhibitor (Olcegepant, BIBN-4096) (**Fig. 6C, D**). In both assays and for both inhibitors we noted a significant reduction of 2D colony-formation and 3D sphere-formation capacity of EwS.

**Figure 6:**
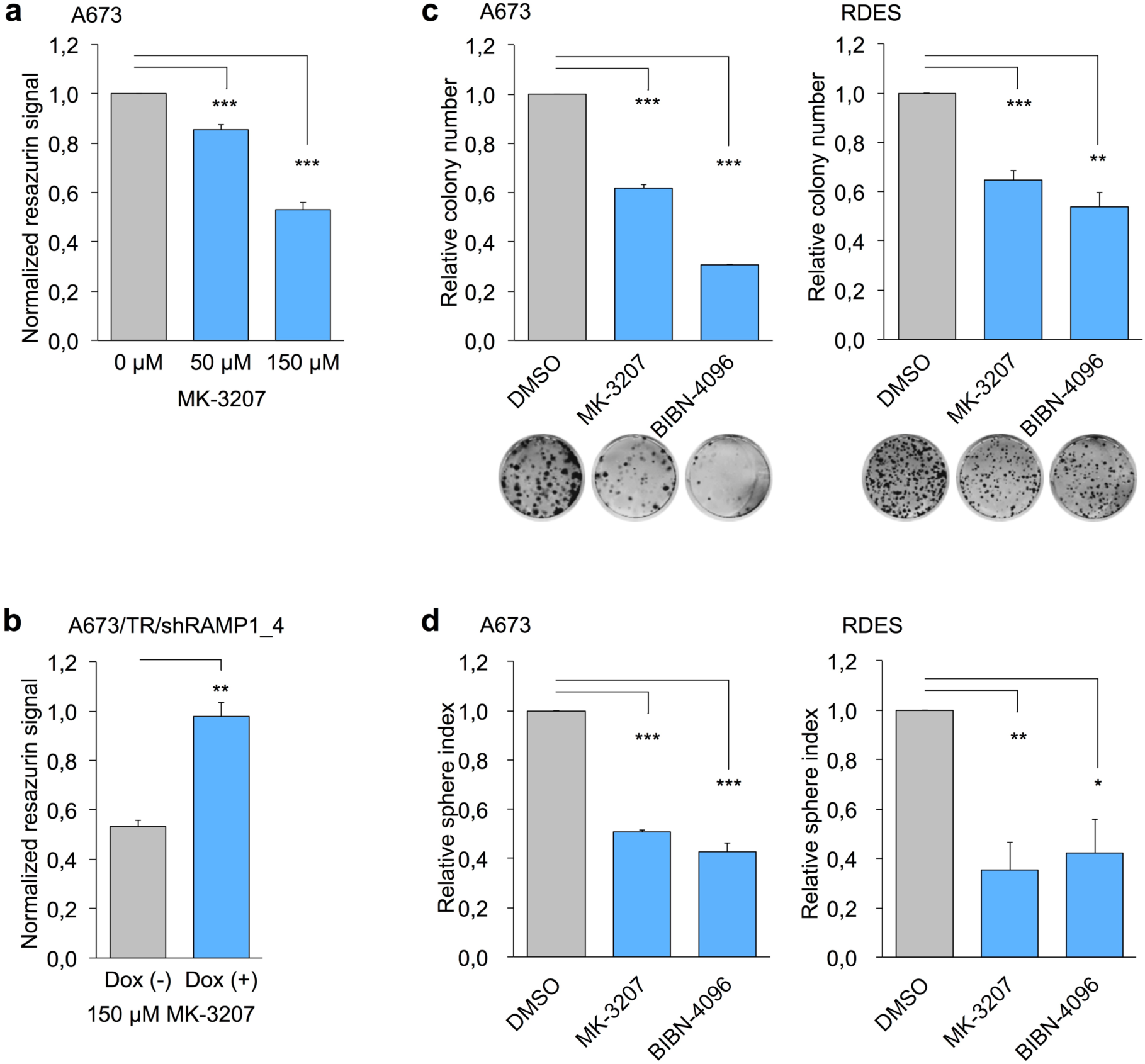
Blockage of the CGRP-receptor by small molecule inhibitors mimics the effect of *CALCB* and *RAMP1* knockdown *in vitro*. **A)** Analysis of cell viability (Normalized Resazurin signal to DMSO control) in A673 EwS cells treated for 72 h with indicated concentrations of MK-3207. The graph shows the dose-dependent relative Resazurin signal reduction. Data are represented as mean and SEM (*n* = 12); unpaired two-tailed student’s t-test. **B)** Comparison of relative Resazurin signal reduction of A673 cells carrying a dox-inducible shRNA against *RAMP1* treated with 150 μM of MK-3207 with/without knockdown of *RAMP1* by additional addition of 1 μg/ml dox to the growth medium. Data are represented as mean and SEM (*n* = 3); unpaired two-tailed student’s t-test. **C)** Analysis of colony forming capacity of A673 (left panel) and RDES (right panel) EwS cells under treatment with the small molecule CGRP-receptor inhibitors MK-3207 (20 μM) or BIBN-4096 (Olcegepant; 100 μM). DMSO served as control for treatment. Representative images of the colonies are shown below. Data are represented as mean and SEM (*n* = 3); unpaired two-tailed student’s t-test. **D)** Analysis of sphere formation capacity of A673 (left panel) and RDES (right panel) EwS cells under treatment with the small molecule CGRP-receptor inhibitors MK-3207 (20 μM) or BIBN-4096 (Olcegepant; 100 μM). DMSO served as control for treatment. Sphere index was calculated by addition of diameter of all existing spheres in one well divided by diameter of spheres in the control well. Data are represented as mean and SEM (*n* = 3); unpaired two-tailed student’s t-test. n.s. *P* > 0.05; * *P ≤* 0.05; ** *P ≤* 0.01; *** *P ≤* 0.001

Together, these data provide further evidence for a functional role of the CALCB/RAMP1-axis in growth of EwS, which could potentially be exploited therapeutically.

## DISCUSSION

Albeit EwS is a disease, which is genetically well characterized, standard therapy still comprises only unspecific cytotoxic approaches. As it is not possible to directly target the underlying cause of the diseases, an abnormal transcription factor, EWSR1-ETS, which is formed by translocation, current strategies to find new and more specific treatment options is to investigate target genes of EWSR1-ETS and determine their potential as possible new therapy targets. To this end we investigated the potential of the CALCB/RAMP1-axis as new target for treatment of EwS and explored its functional role in EwS cells by combining a series of *in situ, in vitro*, and *in vivo* experiments. We found that CALCB is a secreted peptide that shows a highly specific expression pattern among malignant and normal tissues (**Fig. 1A, B, Supplementary Fig. S2**). The high expression of *CALCB* in EwS is likely driven by EWSR1-FLI1 binding to a GGAA-microsatellite at the *CALCB* locus. Since this GGAA-microsatellite is transformed into a *de novo* enhancer upon EWSR1-FLI1 binding but doesn’t show enhancer activity in the absence of EWSR1-FLI1, we speculate that different mechanisms may operate in other tissue types such as trigeminal ganglia to upregulate *CALCB* expression, which remain to be elucidated.

In our long-term knockdown experiments, we observed that silencing of *CALCB* or *RAMP1* reduced growth of EwS cells *in vitro* and *in vivo*. To the best of our knowledge this is the first report of a functional role of CALCB in growth of cancer cells to date. However, we noted a weaker effect of the *CALCB* knockdown as compared to that of *RAMP1* on tumor growth in *in vivo* experiments, which may be caused by residual *CALCB* expression in the EwS cells (around 5% remaining expression), or alternatively by circulating murine Calcb, which might have compensated at least in part for the loss of human CALCB.

In our drug-response assays we found that inhibition of CGRP-receptors with two different small molecules had a similar anti-proliferative effect on EwS cells to that of the *CALCB* or *RAMP1* knockdown. CGRP-receptor inhibitors already showed high efficacy in the treatment of migraine, which is presumably caused via CALCA- or CALCB-mediated vasodilatation in the vicinity of the trigeminal ganglia – the only normal tissue type with physiologically high *CALCB* expression levels found in our analyses (**Fig. 1A**, **Supplementary Fig. S2**). We speculate that repurposing and further optimization of such ‘migraine-drugs’ could perhaps offer novel therapeutic options for EwS patients in the future. As we did not observe differences in density of tumor-associated blood vessels quantified by staining for murine CD31 in immunohistochemistry of our xenograft experiments (**Supplementary Fig. S6**), we assume that the growth-promoting effect of the CALCB/RAMP1-axis is conferred via different mechanisms than one would expect from the vasodilatory effect of CALCB known from the literature^37^.

Collectively, we identified *CALCB* as a highly specifically expressed EWSR1-FLI1 target gene encoding a secreted peptide that promotes growth of EwS cells and show that targeting the CALCB/RAMP1-axis in EwS may offer a new therapeutic approach. Future studies will have to dissect the precise downstream signaling and how the CALCB/RAMP1-axis promotes proliferation of EwS cells, to further explore its therapeutic potential.

## Supporting information

## ACKNOWLEDGEMENTS

We thank A. Sendelhofert and A. Heier for excellent technical assistance, and P. Gilardi-Hebenstreit for technical advice.

## FUNDING

The laboratory of T. G. P. Grünewald is supported by grants from the ‘Verein zur Förderung von Wissenschaft und Forschung an der Medizinischen Fakultät der LMU München (WiFoMed)’, by LMU Munich’s Institutional Strategy LMUexcellent within the framework of the German Excellence Initiative, the ‘Mehr LEBEN für krebskranke Kinder – Bettina-Bräu-Stiftung’, the Walter Schulz Foundation, the Wilhelm Sander-Foundation (2016.167.1), the Friedrich-Baur foundation, the Matthias-Lackas foundation, the Dr. Leopold und Carmen Ellinger foundation, the Gert & Susanna Mayer foundation, the Deutsche Forschungsgemeinschaft (DFG 391665916), and by the German Cancer Aid (DKH-111886 and DKH-70112257). J. Li was supported by a scholarship of the China Scholarship Council (CSC). J. Musa was supported by a scholarship of the ‘Kind-Philipp-Foundation’, M. Dallmayer by a scholarship of the ‘Deutsche Stiftung für Junge Erwachsene mit Krebs’, and T.L.B. Hölting by a scholarship of the German Cancer Aid. M. C. Baldauf, M. F. Orth, and M. M. L. Knott were supported by scholarships of the German National Academic Foundation. M. C. Baldauf was supported by a scholarship of the Max Weber-Program of the State of Bavaria.

## CONFLICT OF INTEREST

The authors declare no conflict of interest.

