## Supplementary material for "Targeting the CALCB/RAMP1-axis inhibits growth of Ewing sarcoma"

### SUPPLEMENTARY INFORMATION

#### Supplementary Table 1

Microarray data accession codes

#### Supplementary Table 2

Mass spectrometric detection of CALCA and CALCB in EwS cell line supernatants

### SUPPLEMENTARY FIGURES

**Supplementary Figure S1** Dallmayer *et al.*

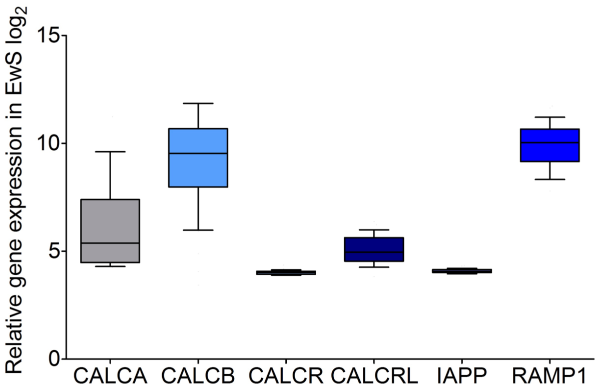


**Supplementary Figure S1: Comparison of *CALCA*, *CALCB*, *CALCR*, *CALCRL*, *IAPP*, and *RAMP1* expression levels in EwS**
Analysis of expression levels of publicly available microarray data of 50 primary EwS tumors are shown in $log_2$-scale. Data are represented as box-plots. Horizontal bars indicate median expression levels, boxes the interquartile range. Whiskers indicate the $10^{th}$ and $90^{th}$ percentile, respectively.

**Supplementary Figure S2** Dallmayer *et al.*

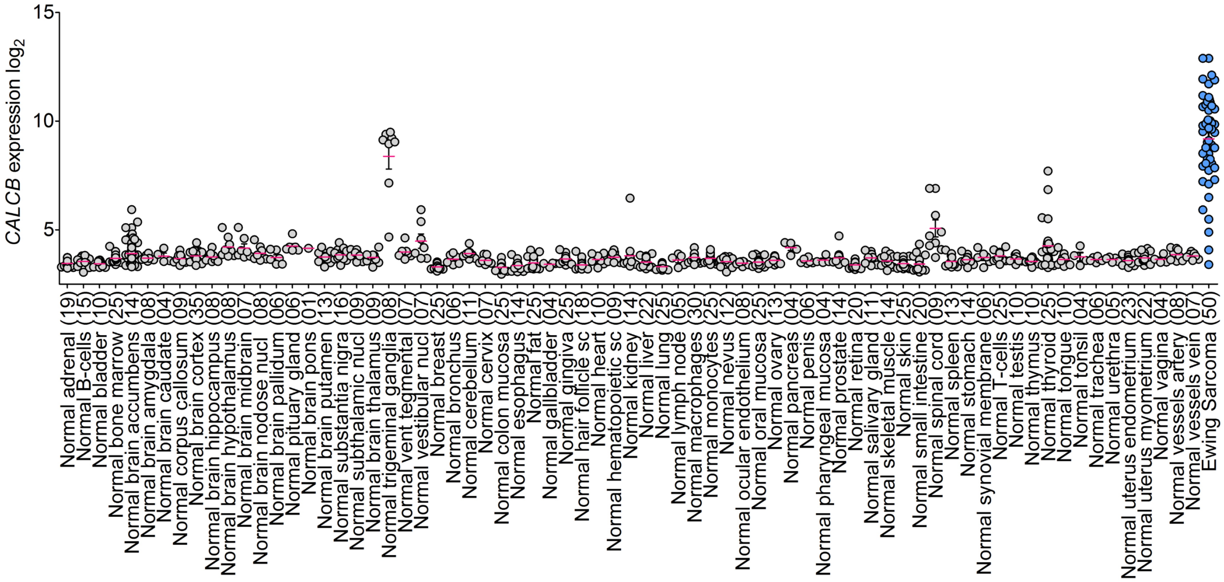


**Supplementary Figure S2: *CALCB* Expression in 71 normal tissue types and EwS**

Publicly available microarray data are represented as scatter plot in $log_2$-scale with mean and SEM. Each dot represents one sample. Number of samples is given in parentheses. EwS is highlighted in blue.

**Supplementary Figure S3** Dallmayer *et al.*

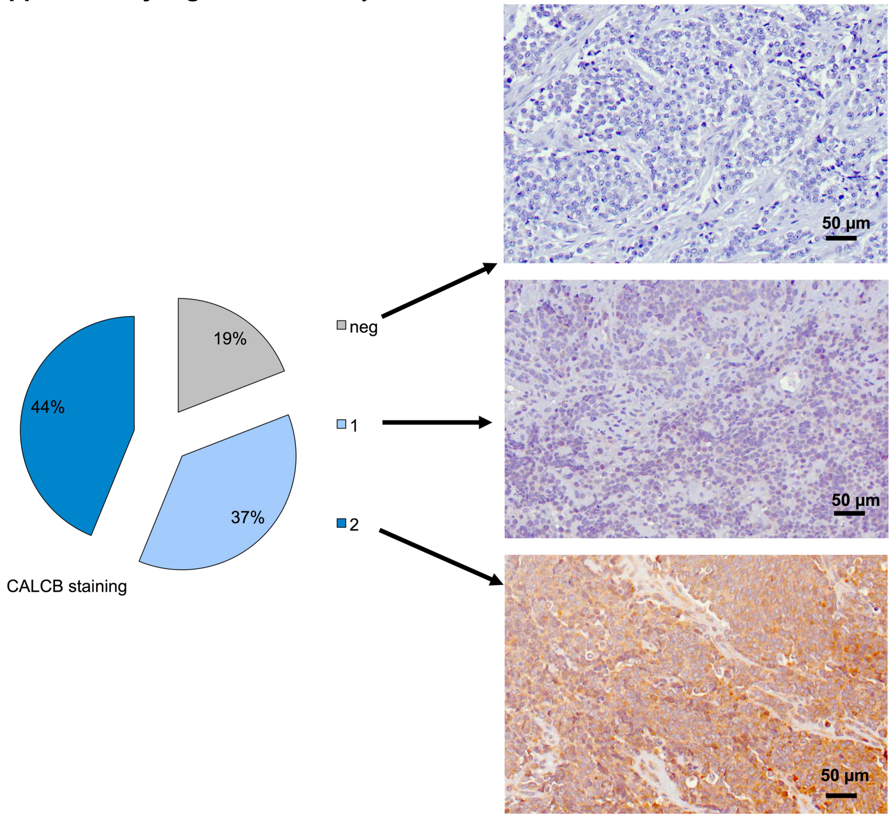


**Supplementary Figure S3: EwS TMA stained for CALCB**
A EwS TMA comprising 89 primary EwS samples was stained with a specific antibody against human CALCB. Micrographs on the right show representative samples with CALCB staining intensity 0 (upper picture), 1 (middle picture), and 2 (lower picture).

**Supplementary Figure S4** Dallmayer *et al.*

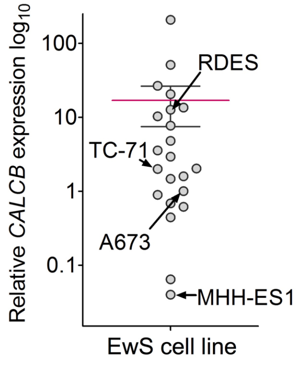


**Supplementary Figure S4: Analysis of relative *CALCB* expression in EwS cell lines**
Relative *CALCB* expression levels of 21 EwS cell lines relative to the A673 EwS cell line were determined by qRT-PCR. Data are shown as scatter plot in $\log_{10}$ scale. Each dot represents the mean *CALCB* expression from two independent replicates for each cell line. Red horizontal bar indicates mean expression, whiskers indicate SEM.

**Supplementary Figure S5** Dallmayer *et al.*

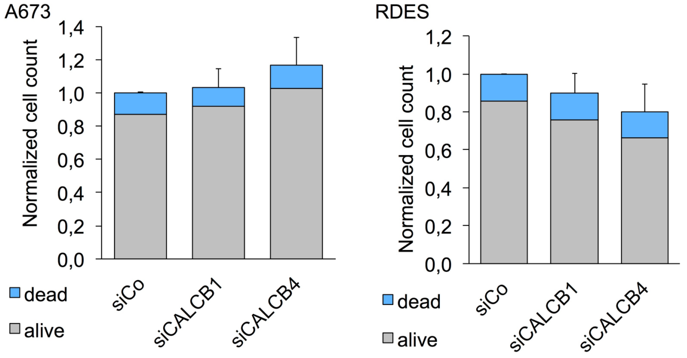


**Supplementary Figure S5: Analysis of short-term proliferation of A673 and RDES EwS cells after silencing of *CALCB***

Cells were transiently transfected with specific siRNAs against *CALCB* (siCALCB1 and siCALCB4) or non-targeting control siRNA (siCo). Cells were harvested after 48-72 h after transfection and the number of viable and dead cells was determined by standardized hemocytometry including cell culture supernatants. Data are represented as mean and SEM ($n_{\text{A673}}$ = 2-4; $n_{\text{RDES}}$ = 3).

**Supplementary Figure S6** Dallmayer *et al.*

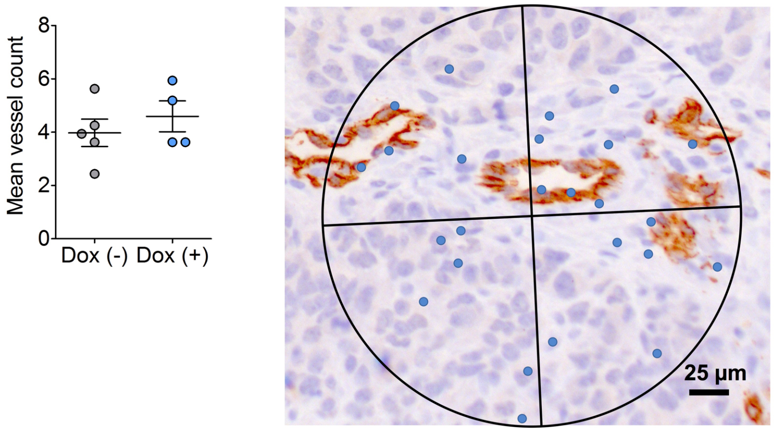


**Supplementary Figure S6: Microvessel density evaluation in CD31 stained xenografts**

Left panel shows mean microvessel count evaluated in murine CD31 stained EwS Xenografts with/without silencing of *CALCB* in the tumors (dox +/–). Each dot represents mean microvessel density evaluated in 4 high power fields of one tumor. Horizontal bars indicates the mean, whiskers the SEM. Right panel: representative CD31 stained slide with Chalkley grid.
